# Ephaptic-Axonal Interactions Predict Radial Biases During Early Neural Self-Organization

**DOI:** 10.1101/2025.11.03.686361

**Authors:** Daniel Rebbin, Casper Hesp, Dimitris Pinotsis, Michael Levin, Wesley Clawson

## Abstract

Communication in the brain seems to not just be mediated by axonal action potential transmission resulting in post-synaptic modulations, but also by electrical field effects of depolarizing neurons, i.e. ephaptic coupling. The significant role of ephaptic coupling in synchronizing neural ensembles has been conjectured to indirectly affect ontogenetic neural circuit development through sub-threshold impacts on spike timing. We therefore hypothesized synchronously firing ensembles to emerge at spatial distances where ephaptic and axonal signals were most temporally correlated. To test our model of ephaptic-axonal interactions during neural self-organization, we compared its predictions to developmental outcomes of cortical rat tissue on high-density multielectrode arrays *in vitro*. We observed a cosinusoidal variation of synchronous activity over radial distances that can be understood to result from the joint effects of ephaptic and axonal signals during gamma-band bursts. Recurrent amplification of this theorized interaction effect during ontogenetic differentiation appears to radially bias the spectral profile of neural activity in the spatial distribution of neural ensembles. While long-term plasticity has been conventionally attributed to synaptic action alone, we show that the interaction effects with ephaptic waves deserve further investigation.

## 1 Introduction

Neuronal networks functioning is known to be constrained through their geometry: Functional coupling depends on, for example, inter-regional distance or gyral curvature. Importantly, this effect runs also in the opposite causal direction: Network geometry *emerges* from functional coupling under resource constraints, where a tradeoff between the functional value of a link and an exponentially increasing wiring costs over distance applies [1]. During developmental symmetry-breaking, axons follow activity-dependent neurochemical gradients and synapses grow between functionally coupled neurons through spike-time dependent plasticity (STDP). In the process of forming sparsely coupled modules, not only are structural connections (network edges) pruned but also neurons themselves (network nodes): Participation in coordinated network activity was found to influence where neurons would survive early apoptosis above and beyond elevated baseline activity [2, 3]. The geometry of where and which neurons settle into a modular network is mediated by neuro-chemical and -electrical signals that guide network self-organization towards a functional attractor configuration.

While axonally mediated synaptic activity has played a prominent role in the investigation of neuronal network formation, endogenous electrical field effects (ephaptic coupling) have received relatively less attention [4]. Although applied electrical fields are known to influence ontogenetic development, guiding axon growth [5] as well as synapse and myelin formation, ephaptic coupling has long been presumed to play at most a secondary role in the differentiation of coupled networks compared to synaptic ‘weight updates’. This is likely due to the weak voltage modulation effect (mostly *<* 1 mV) at endogenous field strengths which is hardly sufficient to push even close neighbors past the firing threshold, for which about 15-20 mV depolarization from rest are required [6, 7]. Dipolar voltage effects have been experimentally well-approximated by a quadratic power law decay with radial distance [7, 8]. Therefore, ephaptic coupling is mostly a subthreshold phenomenon that seems to only have minor effects on single neurons under weak field strengths generated by isolated neuron spiking. This leaves the question: How could ephaptic coupling, despite its weak effect magnitude when viewed in isolation, nonetheless play a decisive role in developmental network formation?

By widening the scale of investigation to the network scale, various studies have shown that weak local field effects can still have strong global effects on network-wide synchrony. Strong dipoles generated by high-synchrony events (bursts) can entrain regions multiple millimeters away [9]. Beyond this, a much simpler mechanism allows for ephaptic coupling effects to in principle reach distant sites far past the reach of the direct dipole field effect that only becomes evident when shifting focus from voltage to *phase* effects. Crucially, power-law decay constrains the field of an *isolated* dipole, not the reach of a spatial phase periodicity in a coupled medium. Therefore, ephaptic coupling can have a non-negligible influence on phase entrainment over large distances even under weak endogenous field strengths. Spatial phase coherence over distance then varies as a standing wave pattern over distance as a function of frequency and signal traveling speed. Ion-flux-mediated ephaptic wave propagation can occur independent from synaptic propagation and has been consistently shown *in vitro* to have discernibly slower traveling velocity (at *≈* 0.1 mm/ms [10–12]) than unmyelinated axonal action potential (AP) propagation (at *≈* 0.6 mm/ms [13, 14]). It has previously been noted that these different propagation mechanisms and velocities are likely to interact in determining spike timing [15]. Thus, the starting point for our experimental rationale is that ephaptic and axonal signals need to be considered *jointly* in investigating STDP at the network scale (Fig. 1a).

**Fig 1.**
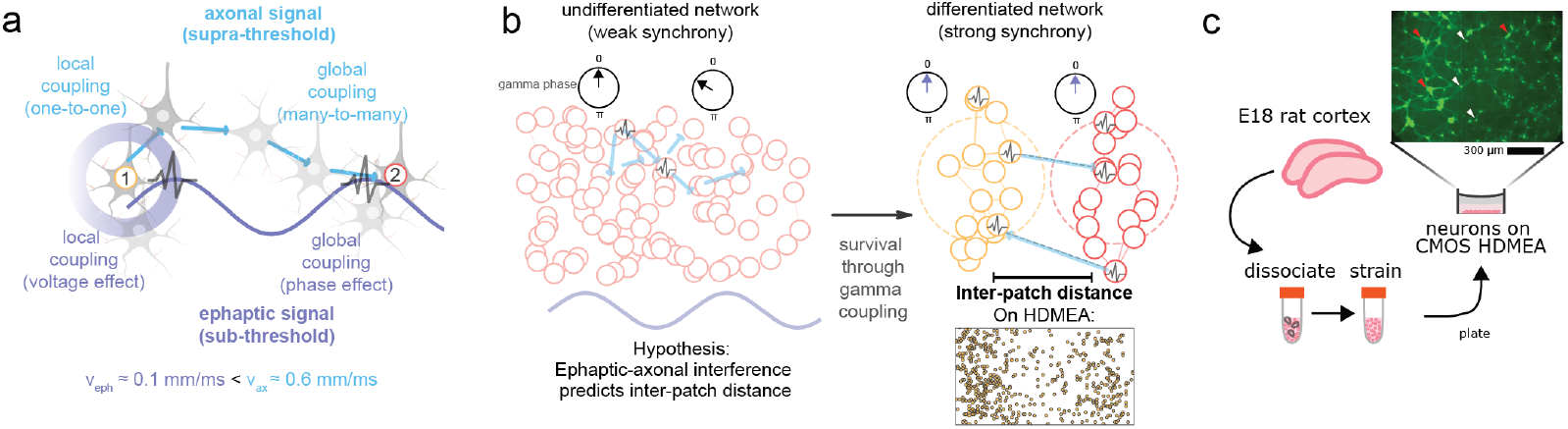
Ephaptic-axonal interference model predicts distance between developing neuronal patches. **a** Field-mediated ephaptic effect propagate with lower speed than axonally transmitted spiking cascades. This difference leads to a global interference effect at a distance determined by the source firing frequency. **b** Coupled patches form during early prenatal development, whose distance are predicted by our ephaptic-axonal interference model. **c** Embryonic E18 rat cortical neurons are plated *in vitro* on a high-density multielectrode array (HD-MEA) to observe spontaneous differentiation dynamics from an isotropic spatial distribution.

Accounting for the temporal cross-correlation between ephaptic and axonal signals, we test in this paper a model for how their velocity difference results in an interference pattern that shows a cosinusoidal periodicity in functional coupling over distance [16]. We hypothesized that these spatial cross-coupling biases during gamma-frequency bursts result in a periodic bias in where we find active neurons in a dissociated culture recorded on high-density multielectrode arrays (HD-MEAs) following network differentiation (Fig. 1b). Within the boundaries of our HD-MEA, where we plated dissociated rat cortical neurons (Fig. 1c), a model that only respects axonal transmission delays (‘axonal-only’) would predict a monotonic (exponential) decay of coupling over distance and an isotropic spatial activity distribution. In contrast, our model (‘ephaptic-axonal’) predicts a periodic deviation from an exponential coupling decay over distance and an anisotropic spatial activity distribution. Under low-gamma-frequency firing, we expect this periodicity to have a wavelength around *≈* 2.5 mm, precisely where the ephaptic-axonal velocity *difference* is expected to drive STDP through coinciding ephaptic excitability shifts and afferent synaptic inputs.

To test this model, we plated dissociated E18 rat cortical neurons on HD-MEAs and recorded spontaneous activity from six wells between DIV 12 and DIV 21. We first measured the two propagation speeds our model requires: supra-threshold spike waves traversed the array at axonal conduction speeds (median 0.62 mm/ms), whereas sub-threshold gamma phase fronts advanced almost an order of magnitude more slowly (0.081 mm/ms), in the ephaptic range. Entering these velocities together with the dominant low-gamma mode (37.76 Hz at DIV 12) into our interference function, we predicted and observed a periodic deviation from the exponential decay of synchronous firing probability over distance, with a wavelength of *≈* 2.5 mm. An unconstrained parameter grid search over a generative model of synchrony-dependent survival then independently recovered this low-gamma regime, reproducing the anisotropic spatial activity distribution that an axonal-only model could not. Together, these results suggest that ephaptic and axonal signals, considered *jointly*, bias activity patterns and early network self-organization.

## 2 Methods

### 2.1 High-Density Multielectrode Array

We used a metal oxide semiconductor (CMOS-based) high-density multielectrode array (HD-MEA) system (“MaxTwo”), produced by MaxWell Biosystems (Zurich, Switzerland), for our local field potential (LFP) recordings. The MaxTwo system records plates of 6 wells (independent HD-MEA arrays), each comprising 26,400 electrodes on a 17.5 µm pitched 3.85 x 2.10 mm^2^ rectangular grid. However, out of those 26400 only 1024 channels can be recorded simultaneously from at a maximum sampling rate of 10 kHz.

Before cells were cultured, the MEAs were pretreated for better adhesion and cell health. Each array was incubated with a 1% Terg-a-zyme-solution (10g/L) at room temperature for 2 hours. Arrays were then washed three times with sterile deionized water. For sterilization, each plate and array were incubated with 70% ethanol for 30 minutes. After, the ethanol was removed and plates washed again with sterile deionized water. Wells were then pre-treated with BrainPhys (STEMCELL Technologies; Cat#05790) and incubated at 37°C, 5% CO2 for 2 days.

### 2.2 Primary Cell Culture

Pre-treated MEAs were washed with sterile deionized water and coated with 0.5mg/ml Poly-d-Lysine (Sigma Aldrich; Cat#A-003-M) in 1x Dulbecco’s Phosphate Buffered Saline with magnesium and calcium overnight (DPBS–). The MEAs were then washed 10 times and airdried before adding 0.02mg/ml Laminin (Millipore Sigma; Cat#L2020) into the NeuroCult device (STEMCELL Technologies; Cat#05713) supplemented with 2 mg/ml L Glutamic Acid, 1M Glucose (final 15mM Glucose), 200mM L Glutamine, and SM1 Neuronal Supplement (STEMCELL Technologies; Cat#05711)) directly on the array. Laminin coated MEAs were kept in tissue culture incubator (37°C, 5% CO2) until cells are plated (1 – 2 hours). E18 Sprague Dawley Rat cortices were obtained from TransnetYX Tissue. The tissue was digested in 6mg/ml Papain in Hibernate (supplied by TransnetYX) and triturated with a silanized Pasteur pipette until completely disassociated. Remaining neurons were counted and resuspended at 2.5 x 106 cells/ml in CPM. 50ul laminin was removed from each MEA and quickly replaced with 1.25 x 105 cells ( 2,500 cell-s/mm2). Cells were maintained in a tissue culture incubator (37°C, 5% CO2). A 90% medium change with CPM is performed on DIV 1. Beginning DIV 5, subsequent medium changes are 50% CMM (BrainPhys with SM1 Neuronal Supplement and 15mM Glucose) every 3-5 days.

### 2.3 Spike detection and recording electrode selection

Spike events were detected from filtered extracellular voltage traces by threshold crossing at 5.25 root-mean-square deviations from the mean, using the peak-amplitude time as the event time. These events were treated as multi-unit activity and were not spike-sorted, because the analyses tested population-level spatial and temporal structure rather than single-neuron identity. Prior to each recording, an activity scan native to the MaxTwo was done to select the electrodes with the highest average spike event rate. For downstream spike analyses, electrodes were ranked by firing rate within each recording and the top active electrodes were retained, with a maximum of 1000 selected electrodes per recording. This procedure was done on each of the days in vitro (DIVs) 12, 14, 21, 22, 23, and 26 for which spontaneous activity in each of the six wells was recorded for 10 minutes.

### 2.4 Burst detection and spike frequency analysis

Instantaneous firing-rate (IFR) traces were computed for each selected electrode from inter-spike intervals and resampled onto a common 50 Hz time grid. Population IFR was defined as the mean across selected electrodes and smoothed with a Gaussian kernel of 0.15 s standard deviation. High-activity epochs were detected using a robust threshold equal to the median population IFR plus three scaled median absolute deviations. Candidate epochs shorter than 30 ms were discarded. This thresholding was used to identify transient network activations without assuming a Gaussian baseline rate distribution.

For burst-level analyses, spike trains were also represented at 1 ms resolution and smoothed with a 3 ms Gaussian kernel. Network participation was evaluated in 10 ms bins. A burst was defined as a high-activity epoch in which at least 5% of selected electrodes participated for at least 20 ms. Burst anchors were assigned to the time of maximal population activity within each burst. IFR samples were then partitioned into low-activity, high-activity non-burst, and burst states. State-specific IFR distributions were summarized by log-spaced histograms and kernel-density estimates, and peaks in the smoothed density were used to identify dominant activity-frequency bands. This family of analyses generated the activity-state and frequency-content panels in Figures 2 and 4. Burst-detection choices follow the principle that MEA burst definitions should combine rate elevation with network participation rather than rely on single-electrode firing alone [17].

**Fig 2.**
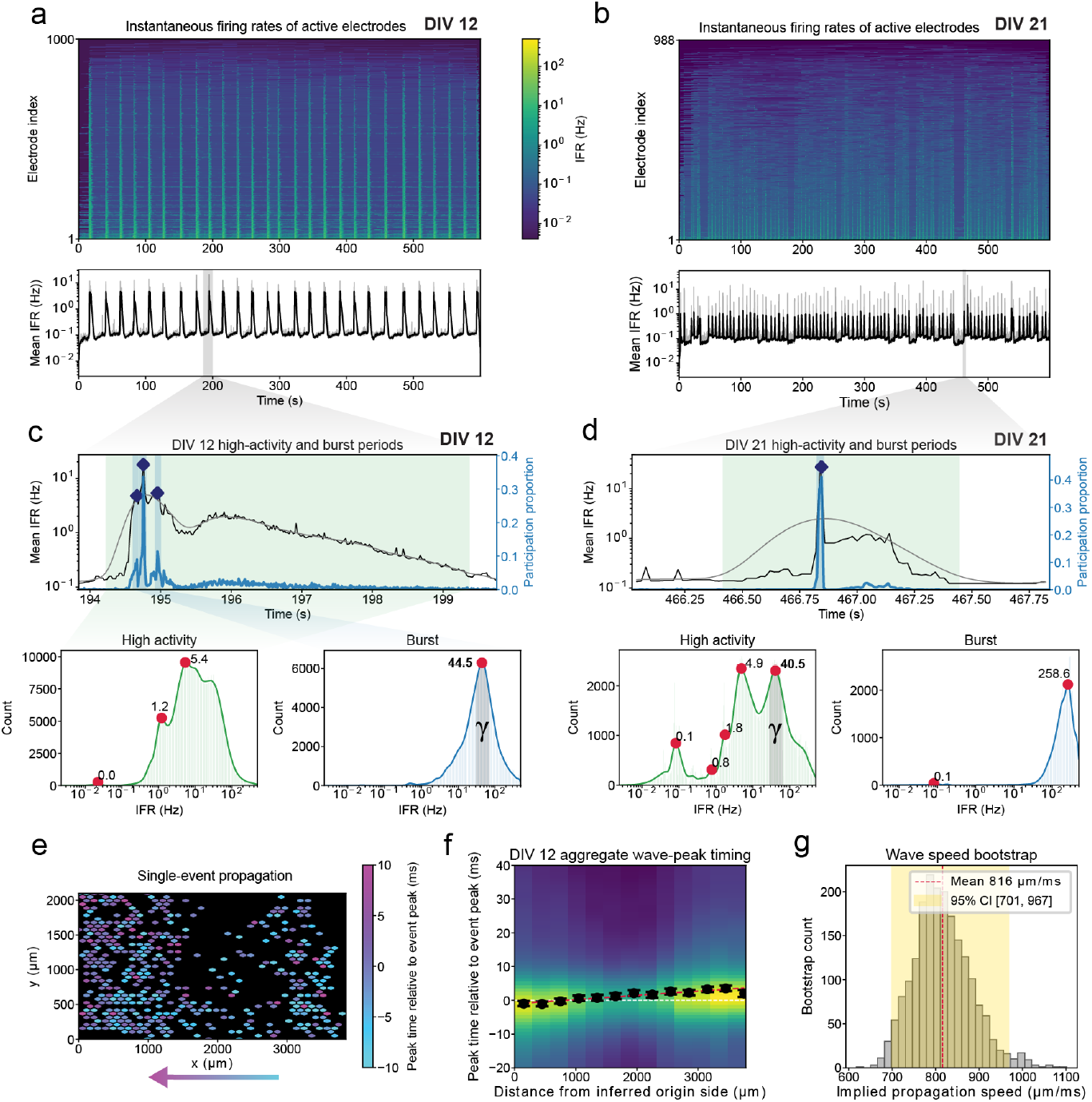
Spike wave propagation during burst episodes. **a** Instantaneous firing rates (IFRs) are recorded for six wells over multiple DIVs across 1000 electrodes each, revealing prolonged high-activity collective states lasting multiple seconds. **b** Same as **a** for DIV 21: More frequent but shorter periods of high-activity are present. **c** Top: activity is categorized into low-activity (white), high-activity (green) and burst (blue) periods. Diamonds indicate peak activity. Bottom: high-activity and burst period IFR histograms are separated and show 30-50 Hz gamma spiking to dominate bursts at DIV 12 (kernel density estimation is used for peak detection). **d** Same as **c** for DIV 21: Low gamma spiking persists during high-activity periods, being led by fast gamma in the 100+ Hz range. **e** Burst events are analyzed individually and show a preference for horizontal propagation (colored according to latency with respect to peak activity) **f** Event-wise burst event origins are detected and their average horizontal propagation speed fit to a spike wavefront (color indicates average IFRs). **g** Bootstrapped horizontal wave velocities are shown to be concordant with axonal transmission speeds under our model.

**Fig 3.**
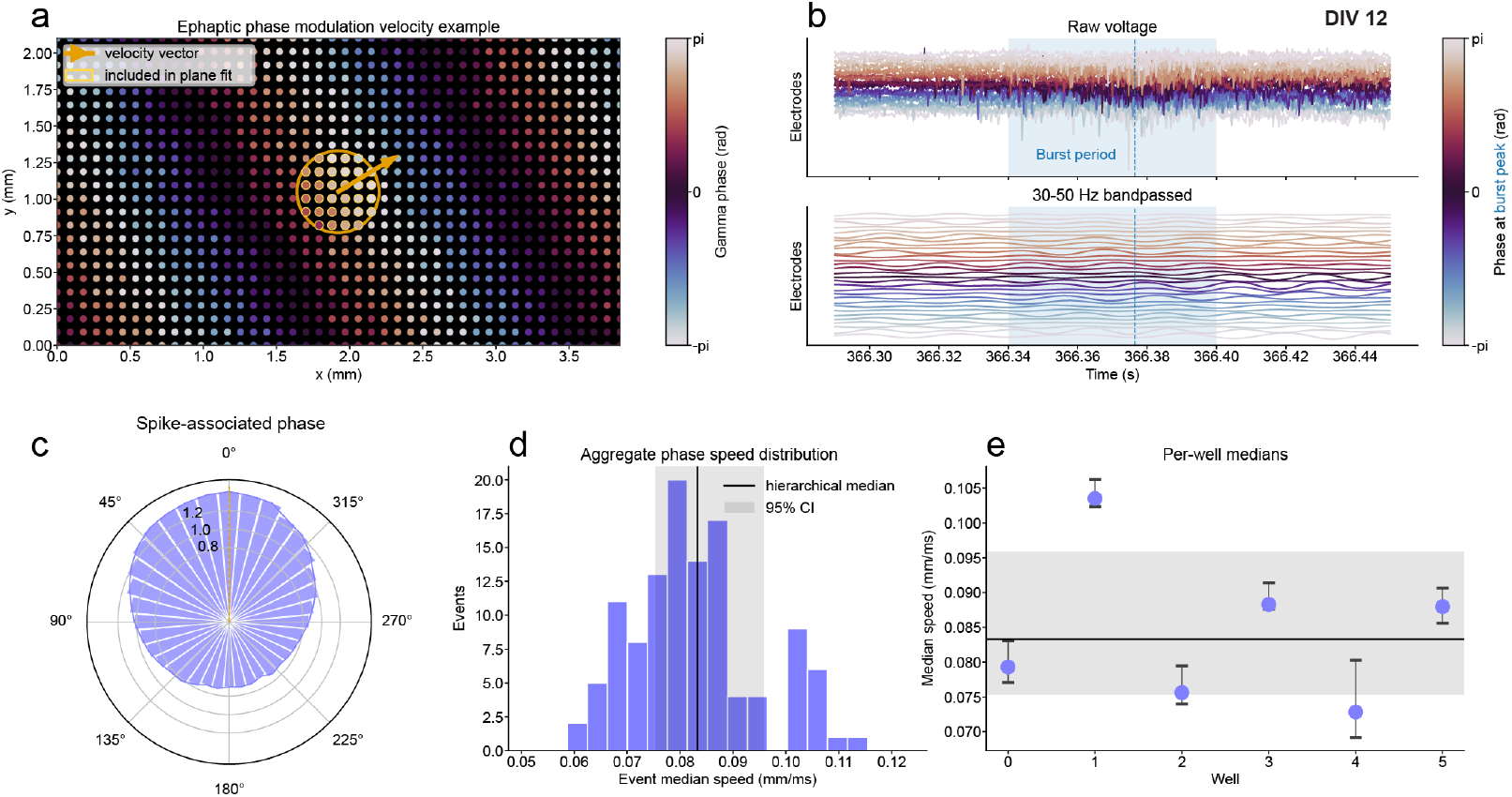
Ephaptic propagation of low-gamma phase velocity around burst events. **a** Method illustration: Phase velocity is determined by a planar fit to a local neighborhood around neurons in their maximally excitable gamma phase (radius = 400 µm). **b** All electrodes (only a subset is shown in the upper panel) receive a 30-50 Hz bandpass filter and are here color-coded with respect to their phase at the burst period peak (vertical blue line). **c** In this low-gamma band, the cultures shows a preference for spiking during a phase of maximum excitability. **d** Median ephaptic phase speeds are aggregated across wells and events are determined to be just below 0.1 mm/ms. **e** Median speed show a strong level of consistency across different wells around 0.084 mm/ms.

### 2.5 Spike-derived propagation speed

For burst propagation analysis, burst-centered spike-density tensors were built from the 1 ms per-electrode IFR traces. Electrodes were binned along the long axis of the MEA in 300 µm spatial bins, requiring at least five electrodes per bin. For each burst and spatial bin, the local activity peak was detected within a predefined window around the burst anchor. Events were classified as left-to-right or right-to-left according to the ordering of peak times across absolute electrode position. Peak times were then re-expressed relative to the event origin, so events propagating in opposite directions could be pooled. Wave speed was estimated by fitting a linear regression line to the mean peak time as a function of distance from burst origin. The inverse absolute slope yielded a speed estimate. Confidence intervals were obtained by bootstrap resampling burst events 2000 times.

### 2.6 LFP phase and event-front analysis

Local field potential analyses used raw HD-MEA LFP recordings from DIV 12 across all six wells. LFP traces were band-pass filtered in the 30–50 Hz band and downsampled to 500 Hz. The analytic signal was obtained with the Hilbert transform, yielding instantaneous phase *θ*_*i*_(*t*) and amplitude *A*_*i*_(*t*) for each electrode *i*.

For each well, an excitable phase *θ*^*\**^ was calibrated from spike-associated LFP phases. Spike phases were assigned to 36 circular phase bins, and phase occupancy was used to normalize spike-phase probability, with a pseudocount of 1 added to each bin. The excitable phase was defined as the bin centre with maximal occupancy-corrected spike probability. Calibration stability was quantified across calibration intervals. In each interval, an independent excitable phase estimate 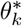 was obtained, and the mean resultant length was computed as

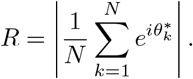

The corresponding circular standard deviation was

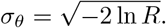

Calibrations were flagged as stable when *R* ≥0.50, indicating that independently estimated excitable phases were concentrated around a common phase.

For event-front analysis, up to 200 burst events per well were scored. For each event, preferred-phase crossings of *θ*^*\**^ were searched from 20 ms before to 40 ms after the burst anchor. Candidate front centres were spaced by the nominal 40 Hz period. For each candidate, the nearest crossing within 8 ms of the candidate centre was assigned as the electrode arrival time. The candidate front closest to the burst anchor was selected, with ties broken by the number of participating electrodes. Events with fewer than 50 assigned electrodes were excluded.

Local propagation was estimated from arrival times. For each electrode, a spatial neighbourhood was formed from electrodes within 0.30 mm; if fewer than 8 neighbours were available, the radius was expanded up to 0.45 mm, with at most 24 neighbours retained. The centre electrode was included in the subsequent fit. Neighbourhoods were excluded when the ratio of the smaller to larger eigenvalue of the local spatial covariance matrix was below 0.10. For each valid centre electrode, neighbouring electrodes were retained only if they had finite arrival times within 8 ms of the centre-electrode arrival. A local arrival-time plane

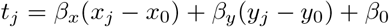

was fit by least squares. Fits required at least 8 neighbours plus the centre electrode, residual RMS *≤* 4 ms, arrival-time span *≥* 1 ms, and pairwise spatial span *≥* 0.20 mm. The local front speed was computed as

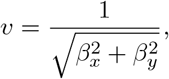

with velocity vector 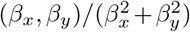. Fits with inferred speed *>* 0.5 mm/ms were treated as unresolved weak-gradient fits and excluded from speed summaries. Each event was summarized by the median speed across quality-controlled local fits. Reported LFP front speeds were summarized by a hierarchical bootstrap that resampled wells and events within wells.

To control for spatially nonspecific front support, spatial nulls were computed from cached event-level arrival tables. For each well, up to 25 events were selected evenly across the event list. For each selected event, 30 null repetitions were generated for each of two nulls: coordinate shuffling, which permuted electrode coordinates while preserving arrival times, and arrival-time shuffling, which permuted arrival times while preserving coordinates. The same neighbourhood construction and local arrival-plane fitting procedure was applied to each null realization. For each well and null type, the 95th percentile of the null distribution of the number of quality-controlled local fits was computed; the final per-well support threshold was the maximum of these two 95th-percentile values. Events were considered front-significant only when their observed number of quality-controlled local fits exceeded this per-well threshold.

### 2.7 Distance-dependent spike synchrony

Pairwise spike synchrony was quantified during burst intervals as excess short-lag co-activity over an interval-jitter null. For each recording, selected electrodes were paired by Euclidean distance and grouped into 100 µm distance bins from 50 to 3500 µm. Spike coincidences were evaluated in a primary lag window of − 2 to 2 ms using 1 ms binned spike matrices. For each distance bin, the observed coincidence probability was compared with surrogate probabilities generated by jittering spikes within burst intervals. The reported excess synchrony was the observed probability minus the mean null probability; bootstrap summaries were computed across recordings and distance bins.

### 2.8 Ephaptic-axonal synchrony model

Following [16], our model parameterized the temporal cross-correlation between two signals emitted at the same time from the same point-like source with different traveling speeds. For simplicity, both signals were assumed to propagate isotropically outwards and propagate at constant velocities, ignoring factors that are known to influence axonal propagation characteristics, such as morphological inhomogeneities [18], axonal arborisation, and variable conduction velocities [14]. In our model, the ephaptic wave travels approximately radially through the tissue volume, supported by the packed arrangement of neurons and glia, rather than strictly along the length of any single axon.

We fit distance-dependent excess synchrony with two nested phenomenological models. The reduced axonal model used an exponential distance dependence,

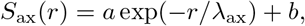

where *r* is inter-electrode distance, *λ*_ax_ is the axonal decay length, and *a, b* are fitted coefficients. The full ephaptic-axonal model added an interference term *C*, so that

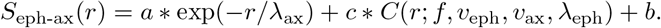

In deriving the temporal cross-correlation (*C*) between our ephaptic and axonal signals, we started by assuming that the source neuron emits with frequency *f* at the same time two signals *s*_*eph*_ and *s*_*ax*_ of the same sinusoidal form sin(2*πft*)

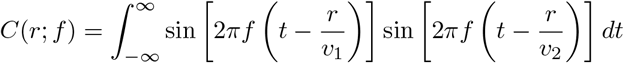

Using the trigonometric identity 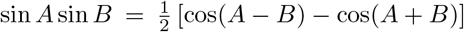 and some algebra, we arrive at the form

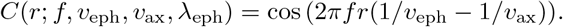

For the Figure 4 fits, *f* was the DIV-specific low-gamma IFR peak, *v*_eph_ was fixed to the DIV 12 LFP front speed, *v*_ax_ was estimated from the median supra-threshold wavespeed across recordings. It is important to emphasise that we do not claim to model microscopic conduction speeds but rather the speed of their emergent effects at the macroscopic scale. The ephaptic coefficient was constrained to be non-negative. Models were fit by nonlinear least squares. Model comparison used residual sums of squares, Bayesian information criterion, and likelihood-ratio tests with one additional degree of freedom for the ephaptic-axonal term.

**Fig 4.**
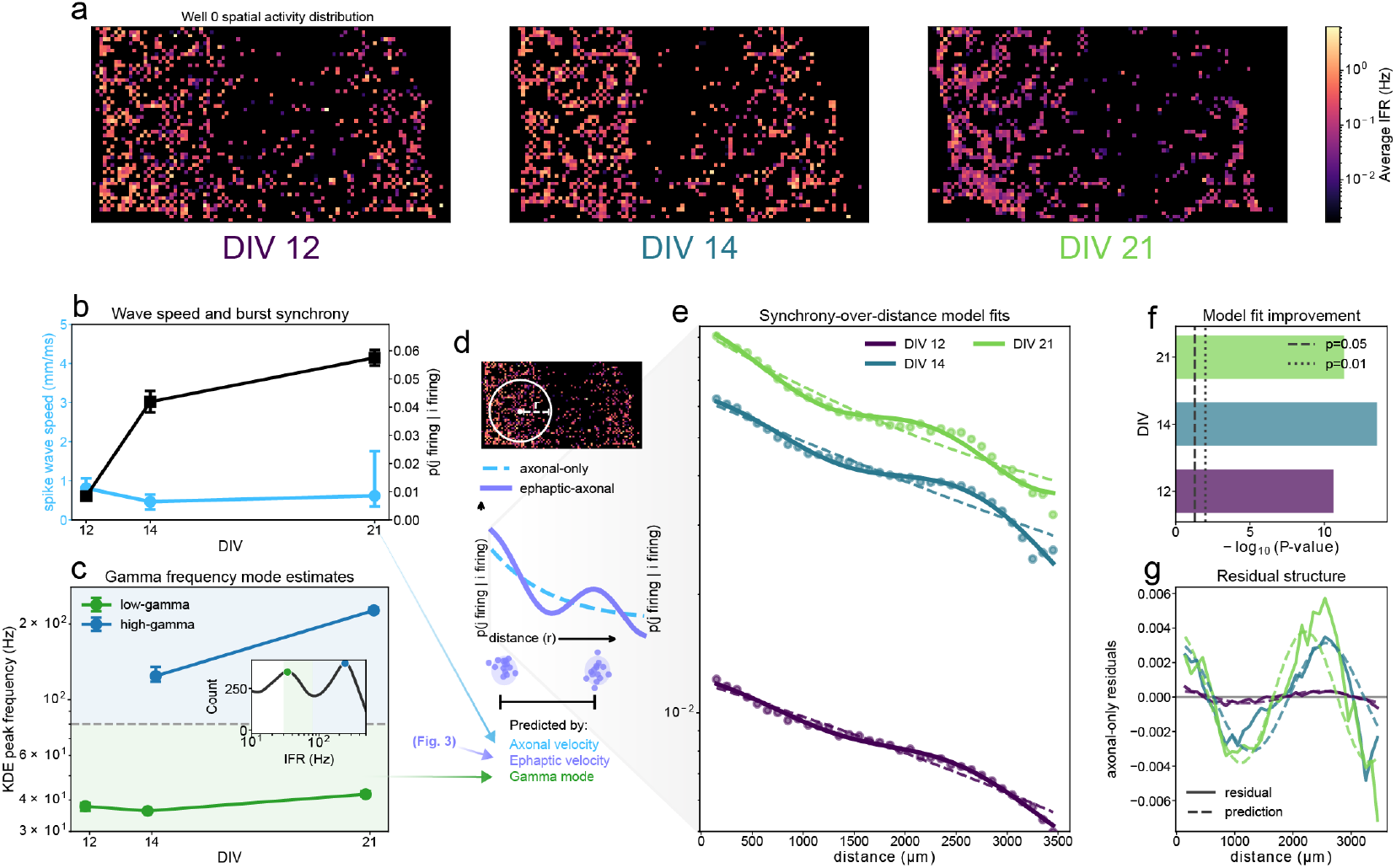
Synchronous firing probabilities over distance deviate from a pure exponential during early DIVs in accordance with ephaptic-axonal interactions. **a** Spatial activity distribution in a single well is shown over DIVs (active electrodes colored according to average IFRs), where the emergence of vertical bands is prominent. **b** Spiking wave speeds are in the axonal conduction speed range but start to diverge from DIV 21 on as spike waves turning into globally synchronized bursts. The average likelihood of synchronous firing above baseline increases over DIVs (95 % CIs shown). **c** KDE peaks (example shown in the inset panel) per recording are separated into low-(30-80 Hz) and high-gamma (*>*80 Hz) and shown over DIVs (95% CIs show average KDE peak frequencies across wells). **d** Low-gamma peaks and axonal (0.6 mm/ms) as well as ephaptic (0.08 mm/ms) propagation velocities yield a characteristic wavelength prediction per DIV. **e** Baseline-corrected increases in synchronous firing probabilities over distance (within *±*2 ms) show clear deviations from an exponential fit, as predicted by our ephaptic-axonal interference model. **f** A likelihood ratio test (LRT) shows significant deviation from an exponential during early DIVs. **g** Residuals from a pure exponential for significant DIVs exhibit a characteristic spatial wavelength.

### 2.9 Spatial differentiation and generative simulations

To test whether the spatial distribution of active electrodes could be explained by ephaptic-axonal interactions, we simulated activity-dependent pruning on an HD-MEA lattice. The Figure 1 schematic simulations used a 110 *×* 60 lattice with 35 µm pitch, corresponding to the HD-MEA downsampled by a factor of two. Initial random plating loss removed 70% of lattice nodes isotropically. The remaining nodes were iteratively pruned until 40% of the remaining nodes were reached. At each update, 20 active nodes were sampled and the half with the lowest interaction scores were removed. In the axonal-only model, the score was the sum of a near-field exponential term and an axonal exponential term. In the ephaptic-axonal model, the axonal exponential term was replaced by the gamma-modulated correlation kernel defined above. This generative spatial-selection model asked whether local activity-dependent survival under different interaction kernels could reproduce the empirical active-electrode imprint.

Figure 5 fit this simulation family to empirical active-site structure on the last DIV analyzed (DIV 21). The empirical target consisted of the top 100 active electrodes from each of six wells. Candidate simulations seeded one virtual neuron per electrode on the HD-MEA (26,400 in total) and randomly eliminated 70% of them to create local inhomogeneities that likely also occur during *in vitro* plating. Out of the remaining nodes, the virtual apoptosis algorithm proceeded until a target empirical electrode count was met. The axonal-only grid varied near-field relative amplitude and axonal decay length. The ephaptic-axonal grid varied near-field relative amplitude and gamma interference frequency. Fit quality was measured with a two-dimensional spatial Kolmogorov–Smirnov statistic comparing empirical and simulated occupancy cumulative distributions over the MEA plane, following multidimensional extensions of the KS distance [19].

**Fig 5.**
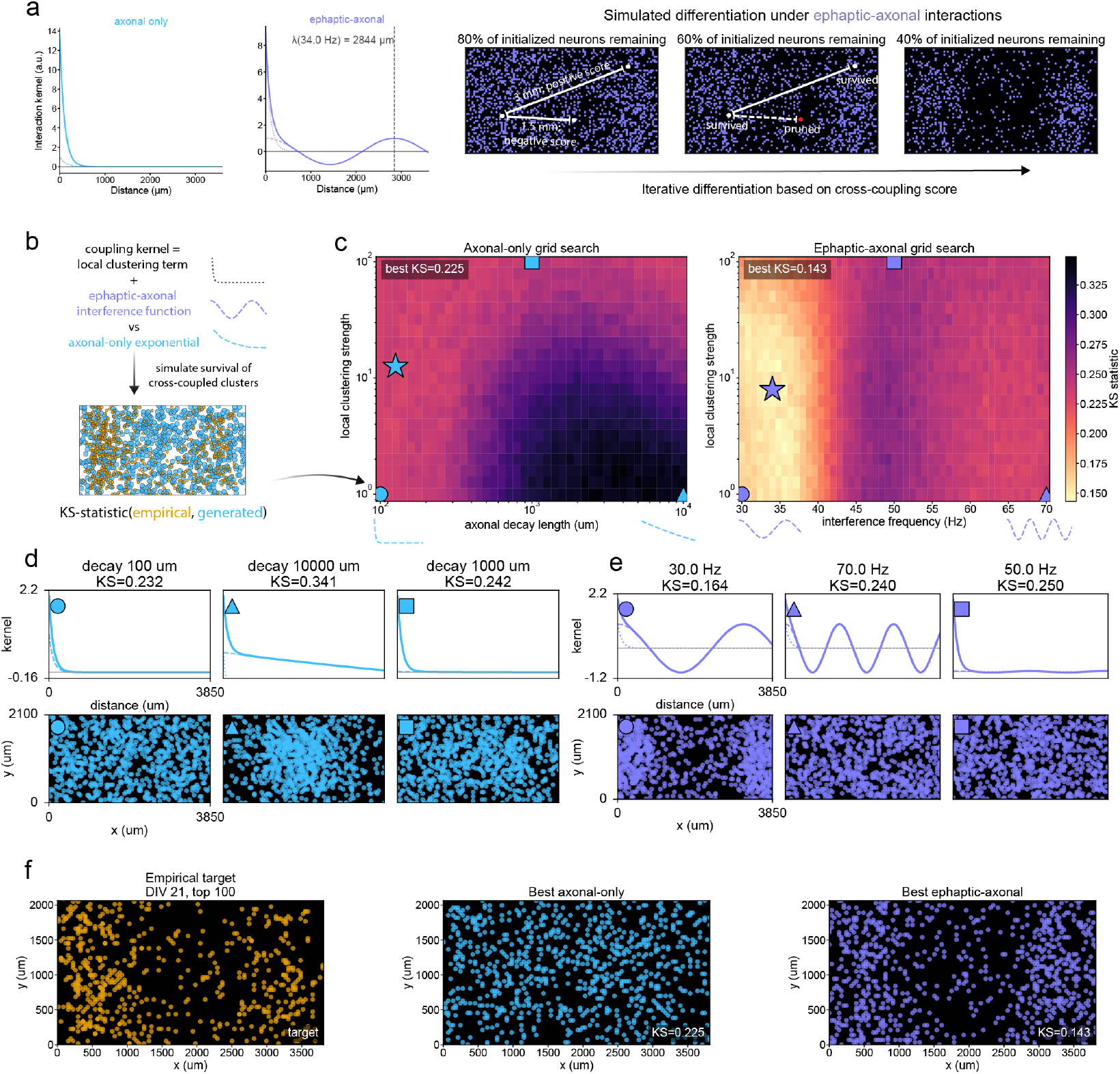
Grid search for best generative model fit to empirical spatial distribution shows superiority of ephaptic-axonal model. **a** Our interference model predicts a synchrony deviation from a pure exponential with a frequency-dependent spatial wavelength. Both models have subsets of neurons being eliminated in an iterative simulation based on the distant-dependent cross-coupling kernels. **b** A local-effect exponential (*λ* = 100 µm) is summed with the coupling kernel from the axonal-only and ephaptic-axonal model, respectively, and used to generate spatial neuron activity distributions. Dis-similarity of different parameter combinations from the empirical distribution at the final DIV recorded is quantified by the Kolmogorov-Smirnov (KS) statistic. **c** Parameter grid search per model shows the lowest KS-statistic for the ephaptic-axonal generative model, specifically for gamma frequency values observed during early DIVs. **d** Different regions of the axonal-only parameter space from **c** are shown with their associated coupling kernels and spatial projections. **e** Same as **d** for the ephaptic-axonal model. **f** The best parameter fit per model shows a close correspondence to the ephaptic-axonal model, while the axonal-only model can only capture local clustering effects.

### 2.10 Statistical reporting and software

Unless otherwise specified, confidence intervals are percentile bootstrap 95% intervals. Bootstrap units were chosen to match the experimental hierarchy: event bootstraps for burst-wave speed, recording or DIV summaries for spike synchrony, and hierarchical well-then-event bootstraps for LFP front speeds. Analyses were implemented in Python using NumPy, SciPy, pandas, h5py and Matplotlib, with project-specific analysis code in the ephax package.

## 3 Results

### 3.1 Low-gamma firing leads the stabilization of burst sequences

In order to understand how ephaptic-axonal interactions could induce structural biases during early network self-organization, we first investigated the presence of low-gamma (30-80 Hz) firing during early network maturation (DIV 12 - 21). For each recording day, we recorded from the 1000 most active electrode sites across the HD-MEA and time-averaged their firing activity to get their spatial distribution. We hypothesized that rhythmic firing in this band would be associated with (1) collective firing events (referred to as ‘bursts’ - see Methods) and (2) the emergence of a structural pattern resolvable within the boundaries of our electrode array. To distinguish frequency modes from spiking activity, we calculated instantaneous firing rates (IFRs), the reciprocal of the inter-spike interval, for the spike traces of the recording electrodes (up to 1000 per well; see Methods). The resulting time series and population traces showed collectively coordinated high- and low-activity states in all wells (n=6) on our first recording day, DIV 12 (Fig. 2a). High-activity periods were often separated by intervals longer than 10 seconds and lasted for multiple seconds (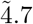seconds on average). For DIV 21 (Fig. 2b), we observed that high-activity periods lasted for only 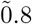seconds, but occurred more frequently (9.6 min^−1^ vs. 2.7 min^−1^ at DIV 12). Spikes became more concentrated in time with less background firing outside of upticks in activity, pointing towards a more stereotyped organization of high-activity periods.

To investigate more closely the role that low-gamma firing played during high-synchrony periods, we needed to distinguish burst states, in which a disproportionately high percentage of electrodes recorded spikes simultaneously across the whole culture, from less synchronous high-activity periods (Fig. 2c). Bursts were indeed dominated by a low-gamma (30-80 Hz) component: multiple bursts with a median duration of 53 ms led a long tail of high activity that decayed only slowly over multiple seconds. At DIV 21 (Fig. 2c), bursts became shorter but increased the proportion of participating sites from 0.1 to 0.25 on average, reflecting increasing levels of synchronous firing. This was accompanied by activity developing into a stereotyped sequence: high-gamma (¿80 Hz) bursts were followed by low-gamma firing, separated by one theta cycle (Fig. 2d). In sum, we observed a nesting of IFR modes that resembles the well-documented theta-gamma nesting [20], with the low-gamma around 40 Hz playing a particularly prominent role during early DIV bursts as we expect under our model.

Our hypothesis requires that low-gamma bursts give rise to propagating spike wavefronts travelling at axonal conduction speeds, since it is the delay between these fast axonal waves and the slow ephaptic field that sets the predicted interference wavelength. Because the HD-MEA resolves propagation most reliably along its long (horizontal) axis, we resolved burst events spatiotemporally and measured, for each electrode, the latency of its peak firing relative to the population burst peak (Fig. 2e). Two features emerged over development. First, a directional polarity preference along this axis sharpened with maturation: at DIV 12 the two propagation directions were near-balanced (56% of events in the more frequent direction), whereas by DIV 14 a dominant direction had emerged (77%; *t*(5) = 3.12, *P* = 0.02, paired *t*-test across wells), and this asymmetry was maintained through DIV 21 (DIV 14 vs. 21: *t*(5) = 0.36, *P* = 0.73). Second, pooling all bursting events across cultures and tracing a ridge of IFRs from the origin edge of the array to the opposite edge (Fig. 2f), we inferred wave speeds of 0.82 mm/ms (95% CI 0.65 − 1.06), 0.46 mm/ms (0.26 − 0.65) and 0.62 mm/ms (0.34 − 1.76) at DIV 12, 14 and 21, respectively (hierarchical bootstrap over wells, 2000 resamples; Fig. 2g). All three estimates lie within the range of unmyelinated axonal conduction speeds reported for comparable cultures *in vitro* ( *≈* 0.4 − 1.0 mm/ms) [13, 14], the wide upper bound at DIV 21 already reflecting the onset of near-instantaneous, globally synchronized bursts. Together, these results establish that low-gamma burst activity propagates as a directed horizontal wavefront at axonal speed, fulfilling a key premise of our ephaptic-axonal interference model.

### 3.2 Gamma excitability wavefronts move with ephaptic propagation speeds

Complementary to supra-threshold spike waves in our ephaptic-axonal interference model, we continued by investigating sub-threshold excitability wavefronts, associated with ephaptic propagation in particular. We therefore used local planar fits to the gamma phase field during burst episodes to estimate ephaptic velocities (Fig. 3a). After applying a 30-50 Hz bandpass filter to the raw LFP data, we anchored local planar fits to gamma phase gradients around peak burst activity for our earliest recording DIV 12 (Fig. 3b). Local arrival-time planes were fit around each electrode, yielding local front direction and speed estimates. Spatial-null correction retained 115 front-significant events across six wells. It is apparent from the inspection of the projection of gamma phases onto the HD-MEA during bursts that spike wavefronts move with a much faster velocity across the array than the excitability fluctuations associated with ephaptic effects that our processing pipeline is aimed at (Supplementary Video XX).

We derived an estimate of spike-associated phases, which were non-uniform in all six wells (Rayleigh’s *R*, all *P <* .001; Fig. 3c). For the example well shown in Fig. 3c, the spike-phase distribution had a mean resultant length of *R* = 0.173 (*P <* .001). Across wells, spike-phase concentration was modest but highly significant (*R* = 0.159 − 0.187, all *P <* .001), as we would expect from ephaptic biases.

The null-corrected event-front speed distribution median was centered at 0.081 mm/ms, with a hierarchical bootstrap 95% confidence interval of 0.077 − 0.101 mm/ms across wells and events (Fig. 3d, e). Per-well median speeds were broadly consistent with the aggregate estimate, although one well’s median (0.105 mm/ms) exceeded the aggregate. These propagation speeds are far below classical axonal conduction velocities and are consistent with a slowly propagating ephaptic recruitment process (*≈* 0.1 mm/ms).

### 3.3 Emerging inter-patch distances correspond to long-range periodicity in synchrony

Having obtained empirical estimates of the different propagation speeds of axonal and ephaptic effects, we investigated whether low-gamma frequency oscillations would predict the wavelength associated with the horizontal trough in activity across the horizontal dimension of our cultured networks. For most of our wells, the characteristic horizontal anisotropy had already started to emerge at DIV 12 and remained stable over DIVs (Fig. 4a).

In order to assess whether our ephaptic-axonal interference function predicts deviations in synchrony that could have led to these inter-patch distances along the horizontal dimension, three key parameters are required to determine the spatial wavelength: axonal and ephaptic wave propagation speeds as well as the dominant low-gamma mode. These jointly modulate the radial distance at which synchrony is expected to drive synchronous cluster formation, where fast and slow signals are expected to fall back into phase (Fig. 4d).

Although there were some fluctuations over DIVs in the speed with which spiking waves horizontally traversed the electrode array, the vast majority consistently reflected propagation speeds in the axonal conduction speed range with a median of 0.62 mm/ms (95% CI: 0.40 − 0.97 mm/ms) (Fig. 4b). These speeds only tended to go towards the supra-axonal speed range (*>* 1 mm/ms) once wave travel time delays vanished in globally synchronized bursts. The likelihood that any two sites fired synchronously (recording a spike within *±*2 ms of one another above expected baseline coincidence rates, see Methods) increased significantly from 0.8 to 5.8 percent from DIV 12 to DIV 21 (*P <* 0.001). Hence, we see an increasing recruitment of synchronously firing neurons over DIVs, while the supra-threshold waves accompanying their synchronization are likely generated through axonally conducted spiking cascades.

In addition to the conduction velocities estimated, the stable presence of a low-gamma mode is required to determine the spatial wavelength. Separating the low-(30−80 Hz) from high-gamma KDE peaks (*>* 80 Hz) revealed a temporally stable low-gamma mode estimate of 37.76 Hz at DIV 12 (36.18 Hz for DIV 14, 42.14 Hz for DIV 21) (Fig. 4c) that entered our wavelength prediction. Given this set of parameters, we fitted two models to the synchrony levels over radial distance from spike origin site: The axonal-only model received an inverse exponential fit per DIV, while the ephaptic-axonal model added to that an extra term for the interference parameter with our model-specific radial wavelength (Fig. 4d). Our hypothesis suggested that synchronous firing probabilities should have cosine residuals from a pure exponential with a radial wavelength between 2 and 3 mm for the low gamma modes around 40 Hz detected. In accordance with this prediction, we observed over DIVs a more pronounced deviation in synchrony-over-distance from a pure exponential drop-off (Fig. 4e, axonal-only residuals isolated in Fig. 4g). For each of the DIVs, the improvement in fit afforded by the ephaptic interference term was substantial, reducing the residual sum of squares by 73.0%, 81.7% and 75.5% at DIV 12, 14 and 21, respectively. Adding this single term significantly improved the fit over the axonal-only model at every DIV (likelihood-ratio test, df = 1; DIV 12: *χ*^2^ = 44.6, *P <* .001; DIV 14: *χ*^2^ = 57.8, *P <* .001; DIV 21: *χ*^2^ = 47.8, *P <* .001) and was decisively favored by the Bayesian information criterion (ΔBIC = − 41.0, −54.3 and −44.2, respectively) (Fig. 4f). The interference term predicts radial wavelengths of *λ* ≈ 2.5, 2.6 and 2.2 mm at DIV 12, 14 and 21. Consistent with this, the residual of the axonal-only fit peaked at 2.5−2.7 mm across DIVs (Fig. 4g), matching the predicted wavelengths to within one to three 100 µm distance bins. We conclude that an important degree of freedom in spike synchrony over distance is consistent wity our ephaptic-axonal interference model.

### 3.4 Low-gamma ephaptic-axonal interference reproduces spatial anisotropy in a generative model

After detecting significant deviations from a pure exponential drop-off from the source, we sought to translate these functional insights to a spatial differentiation patterns through the simulated pruning of the least synchronized neurons. This directly maps functional synchrony onto geometrically constrained structure of the developing network. After a simulated ‘plating’ of virtual point-like neurons on the HD-MEA under spatial homogeneity, our algorithm proceeded iteratively in the following way: A subset of neurons are randomly selected and each scored by the distance- and model-dependent synchrony contributions of their neighbors. Under the assumption that these coupling contributions define their survival likelihood during early apoptosis, the lowest-ranking half is iteratively eliminated until the count of remaining neurons matches the empirical density at the final DIV analyzed. We pitted the axonal-only model’s exponential against the cosine function from our ephaptic-axonal interference model, each additionally receiving a positive local clustering term (with a fixed decay length of 100 µm to account for direct neuronal coupling). (Fig. 5a). The spatial pattern thus generated was then compared against the empirically most active electrode location with the Kolmogorov-Smirnov statistic (Fig. 5b).

Across the parameter grid, the ephaptic-axonal model reached a markedly closer correspondence to the empirical active-site distribution than the axonal-only model (Fig. 5c). The best axonal-only configuration plateaued at a Kolmogorov-Smirnov distance of *KS* = 0.225 (axonal decay length*≈* 126 µm), whereas the ephaptic-axonal model attained *KS* = 0.143 at its optimum. This optimum occurred at an interference frequency of 34 Hz, squarely within the low-gamma band that dominated early-DIV bursts (cf. the 37.76 Hz mode at DIV 12), with the lowest *KS* values confined to the *∼*31− 36 Hz range (Fig. 5c). Notably, this optimum was reached without constraining the frequency to the measured value, so the grid search independently recovered the low-gamma regime (34 Hz vs the measured 37.76 Hz), both predicting *λ≈* 2.8 mm within our 2–3 mm hypothesis window.

Inspecting the coupling kernels underlying each regime clarified the source of this difference: the axonal-only exponential could only seed local clustering and therefore generated a macroscopically isotropic field, whereas the gamma-tuned interference kernel reproduced both the characteristic inter-patch spacing and the horizontal anisotropy of the empirical imprint (Fig. 5d-f). Because the predicted wavelength (*≈* 2.8 mm) approaches the array’s long dimension but exceeds its short dimension, the radially symmetric interference expresses as banding along the horizontal axis. In this way, the rectangular geometry acts as a symmetry-breaking constraint. The generative simulation thus recapitulates, at the level of spatial structure, the same low-gamma ephaptic-axonal interaction that accounted for the synchrony-over-distance deviations, indicating that recurrent activity-dependent survival under this interference kernel is sufficient to bias an initially homogeneous network toward its observed radial organization.

## 4 Discussion

Our results suggest that ephaptic-axonal interference contributes to how neuronal networks self-organize into spatially segregated clusters. Specifically, we captured their almost order-of-magnitude difference in conduction velocities of low-gamma wave signals and demonstrated how they can jointly explain the radial distance at which different patches emerge from a homogeneous seed distribution. We hypothesized that periodic deviation from an exponential synchronous firing probability decay over distance would be reflected in the geometric structure and measured its wavelength to be in close agreement to our prediction. Building on the evidence that synchronous clusters are more likely to survive early apoptosis and network differentiation [2, 3] in a generative model, our model translates the presence of low-gamma firing dominance during early network burst events onto the geometric structure of dissociated cultures *in vitro*. The measured low-gamma frequency was independently recovered from a parameter grid search only under the ephaptic-axonal model assumptions. Thus, neither axonal wiring nor ephaptic coupling alone accounts for the observed patterning; only their interaction at the macroscopic scale does.

It is important to emphasize the macroscopic nature of our proposed ephaptic-axonal interference mechanism does not stand in contradiction with local ephaptic and axonal effects acting on much shorter scales than the wavelength of multiple millimeters we observed. Instead, finite traveling speeds of such local effects in a coupled network induce a periodic phase field with wavelengths independent from their local reach [21, 22]. The insufficiently unpacked distinction between immediate voltage effects and collective phase effects has likely underwritten the belief that ephaptic effects are likely negligible beyond the immediate vicinity of single neurons [4]. Biophysical simulations have shown that the ephaptic voltage modulation effect from a single depolarizing neurons is usually too weak to elicit action potentials in immediately neighboring neurons by itself [6]. However, our model proposes a mechanism how these subtle effects can affect synchrony and, by extension, developmental patterning at much larger scales: Ephaptic effects induce a subtle phase shift in excitability that biases the likelihood that afferent signals elicit synchronous action potentials [23, 24]. This is consistent with our observation show only a weak, yet significant, preference for spiking during the globally most excitable gamma phase. Recurrent amplification of this bias over burst cycles makes it much more likely to entrain distant neurons in an excitable phase [25, 26]. This represents a network-level effect, whose periodicity in radial synchrony and spatial patterning is predicted by our ephaptic-axonal interference model.

Previous literature has mostly considered the geometric properties of cortical networks to be the results of synaptic weight updates, where axonal wiring probabilities decay exponentially over distance [1, 27]. It is important to note that a model which only considers axonal conduction delays also predicts periodically varying functional coupling over distance, when considering emergent phase structure in neurons as coupled oscillators. However, given the observed low-gamma firing at early DIVs, the implied wavelength (0.6 mm/ms / 40 Hz = 15 mm) for a stable configuration far exceeds the dimensions of our HD-MEA (4.4 mm diagonal length) due to high axonal conduction velocities. Therefore, the axonal-only model falls back to an inverse exponential in wiring probability over distance, for which even a double-exponential for long- and short-range connections [28] could only reproduce local clustering (below *≈* 500 µm) but not the macroscopic radial periodicity at the millimeter-scale we observed in our cultures. Accounting for ephaptic biases to phase fields beyond axonal wiring graphs would explain traveling wave speeds in the ephaptic conduction range, which have often been attributed to mean-field effects from axonal cascades [29, 30].

Our propagation speed measurements are derived from suprathreshold spiking waves and subthreshold phase gradients under mean-field assumptions. It could be argued as a limitation to our approach that we did not measure axonal and ephaptic propagation speeds directly but rather the macroscopic effect of their wave-like propagation at the collective scale. The choice to model both signals as waves propagating with constant speed through homogeneous tissue under mean-field assumptions was done intentionally, though. Under this assumption, local inhomogeneities in axonal branching and field gradients average out, sacrificing biophysical detail for a focus on collective-scale effects. It is important to note that the two different wave speeds we measured are nonetheless in agreement with axonal and ephaptic propagation speeds, respectively. The observed median local phase propagation speed of 0.08 mm/ms, almost an order of magnitude below axonal conduction speeds, appears distinctly indicative of ephaptic coupling [8, 10, 31, 32].

Another interesting observation from our data, which also is crucial to our ephaptic-axonal wavelength prediction, is the prevalence of low-gamma firing rates during bursts by DIV 12, when a majority of synchrony-dependent apoptosis has happened [33]. Dissociated neuron *in vitro* form a giant connected component at DIV 4 already [34], show spontaneous synchronized activity around DIV 4-6 [35], self-organizing towards stereotyped burst that we also observed by DIV 12 [36]. The gradual maturation of GABA-ergic interneurons seems to be essential for driving this large-scale synchronisation [37]. Somatostatin (Sst) interneurons have their preferred firing frequency around between 30 and 50 Hz [38], orchestrating gamma-theta cycles [20], and mature earlier than Paravalbumin (Pvalb) interneurons, which more frequently engage in fast-gamma ripples (150-250 Hz) [39, 40]. These observations ground our finding of low-gamma dominance during early DIVs in known stages of functional cell differentiation. Complementary to this interpretation, Sst interneurons have recently been shown to preferentially entrain to weak ephaptic fields in the low-gamma range whereas Pvalb interneurons require fast-gamma oscillations around 150 Hz [41]. Hence, the particular role that SST interneurons play in generating the activity-dependent survival patterns we seek to explain deserves further investigation.

Translating these insights from our effectively two-dimensional dissociated cultures *in vitro* onto three-dimensionally embedded brains *in vivo* and *in utero*, we speculate that the proposed mechanism should play a particularly important role in pattern formation in superficial layers, specifically L2 and L3, where gamma-band firing is most prominent [42]. L2 and L3 are the layers with the highest proportion of unmyelinated horizontal connections between cortical columns [43], something we would expect under our model, given the predominance of low-gamma firing in those layers [44]. The fact that no visual experience is needed to drive this pattern [45] points towards an inherent prenatal pattern formation bias that the effect we propose might help explaining.

A key limitation of this study is that we relied exclusively on observational outcomes from spontaneous activity and thus cannot finally rule out confounds, such as millimeter-scale plating anisotropies, to our explanation of spatial activity distribution. However, our model makes very specific predictions about the cosine-like shape of residuals from exponential decay over distances and the associated wavelength. Crucial here seems to be the timing of the stimulation during sensitive periods of early development: Apoptosis rates are highest before DIV 14 [33], when functional network roles have already become “locked in” [46]. Therefore, inducing oscillations at the lower and higher end of the low-gamma spectrum around DIV 8-10 should lead to noticeably different structural outcomes according to our generative model. Alternatively, changing the network’s plating dimensions is predicted to have a similar effect in producing spatial periodicity in structural outcomes. Modulating in future experiments firing frequency and geometry beyond those recorded here will be crucial to test the robustness of our model.

Taken together, our results suggest that the spatial self-organization of developing neuronal networks is shaped not by axonal wiring alone, but by its interference with slower, field-mediated ephaptic signals during low-gamma bursts. By converting the dominant firing frequency of early bursts into a millimeter-scale spatial wavelength, this mechanism offers a macroscopic, collective-scale account of how an initially homogeneous network acquires a periodic structural bias. Therefore, ephaptic coupling seems to play a globally constraining role in the self-organization of neuronal networks that future studies should jointly investigate with axonally transmitted action potentials they emanate from.

## Acknowledgements

This material is based upon work supported by the Air Force Office of Scientific Research under award number FA9550-22-1-0465. Any opinions, findings, and conclusions or recommendations expressed in this material are those of the author(s) and do not necessarily reflect the views of the United States Air Force. D. R. gratefully acknowledges financial support through the Amsterdam Science Talent Scholarship and the German Scholarship Foundation.

## Declarations

The authors declare no competing interests.

